# Loss of CTLH component MAEA impairs DNA repair and replication and leads to developmental delay

**DOI:** 10.1101/2025.07.23.666279

**Authors:** Soren H. Hough, Satpal S. Jhujh, Samah W. Awwad, Simon Lam, John C. Thomas, Oliver Lewis, Thorsten Mosler, Aldo S. Bader, Lauren E. Bartik, Shane McKee, Shivarajan M. Amudhavalli, Estelle Colin, Nadirah Damseh, Emma Clement, Pilar Cacheiro, Anirban Majumdar, Damian Smedley, Isabelle Thiffault, Guido Zagnoli Vieira, Rimma Belotserkovskaya, Stephen J. Smerdon, Petra Beli, Yaron Galanty, Christopher J. Carnie, Grant S. Stewart, Stephen P. Jackson

## Abstract

Ubiquitin E3 ligases play crucial roles in the DNA damage response (DDR) by modulating the turnover, localization, activation, and interactions of DDR and DNA replication proteins. To gain further insight into how the ubiquitin system regulates the DDR, we performed a CRISPR-Cas9 knockout screen focused on E3 ligases and related proteins with the DNA topoisomerase I inhibitor, camptothecin. This uncovered the CTLH ubiquitin E3 ligase complex — and particularly one of its core subunits, MAEA — as a critical regulator of the cellular response to single-ended DNA double-strand breaks (seDSBs) and replication stress. In tandem, we identified patients with variants in *MAEA* who present with neurodevelopmental deficits including global developmental delay, dysmorphic facial features, brain abnormalities, intellectual disability, and abnormal movement. Analysis of patient-derived cell lines and mutation modeling reveal an underlying defect in HR-dependent DNA repair and replication fork restart as a likely cause of disease. We propose that MAEA dysfunction hinders DNA repair by reducing the efficiency of RAD51 loading at sites of DNA damage, which compromises genome integrity and cell division during development.

## Introduction

As DNA is constantly attacked by exogenous and endogenous agents, DNA repair pathways must function efficiently to avoid accumulation of deleterious genomic alterations^1^. There is much crosstalk between the DNA replication and DNA repair processes, exemplified by RAD51, which is essential for homologous recombination (HR)-dependent repair of DSBs and protection/restart of damaged replication forks^2–5^. Consequently, germline loss-of-function or hypomorphic mutations in DNA damage response (DDR) genes often compromise both DNA repair and replication, leading to clinical phenotypes including neurodegeneration, immunodeficiency, skeletal abnormalities, intellectual dysfunction, bone marrow failure, and growth delay^1,5^.

Protein ubiquitylation is critical in promoting the recruitment and retention of proteins to DNA damage sites, regulating DNA repair and replication protein turnover, and/or altering their enzymatic activities^6–8^. Ubiquitylation in human cells is tightly regulated and mediated via enzymatic cascades involving either of two E1 activating enzymes, one of ∼40 E2 conjugating enzymes, and over 600 known E3 ligases, and various associated factors such as substrate adaptors that enhance target specificity^6,9,10^. Multiple E3 ubiquitin ligases, including those involved in the DDR, have previously been linked to inherited and *de novo* pathologies in humans associated with diverse clinical phenotypes. These include immunodeficiency and neurodegeneration (e.g., RNF168), growth delay and cancer predisposition (e.g., BRCA1), neurodevelopmental abnormalities (e.g., RFWD3), and primordial dwarfism (e.g., TRAIP)^10–15^.

Here, using CRISPR-Cas9 screening^16^, we identify the multi-subunit CTLH (C-terminal to LisH) ubiquitin E3 ligase complex as a regulator of DNA repair and replication. We demonstrate that CTLH — and specifically its RING domain-containing MAEA subunit required for catalytic activity — promotes RAD51 loading at DNA damage sites. We also show that loss of MAEA function severely impairs HR, as well as replication fork progression, protection, and restart. Furthermore, we describe a cohort of patients with pathogenic *MAEA* variants that exhibit neurodevelopmental defects and abnormalities in the cellular replication stress response. This highlights the importance of the CTLH complex in preventing disease driven by genome instability.

## Results

### Loss of the CTLH complex sensitises cells to seDSB inducing agents

To identify ubiquitylation components mediating cellular responses to single-ended DNA DSBs (seDSB), we generated a focused single-guide RNA (sgRNA) CRISPR-Cas9 knockout library targeting 886 E3 ligases and associated substrate adaptors and used this to screen for genes that affect sensitivity of U2OS cells to the topisomerase I (TOP1) inhibitor, camptothecin (**Fig. 1A**, **Supplemental Table 4**). Camptothecin-induced seDSBs are S phase specific and repaired via HR^17^; thus, the screen identified E3 ubiquitin ligases with known roles in HR, including RNF168, BRCA1, and BARD1 (**Fig. 1B**). The CRISPR screen also identified subunits of the CTLH complex as conferring resistance to camptothecin. These included MAEA, WDR26, RMND5A, and GID8, and RANBP9 (**Fig. 1B, Supp. Table 4**), which represented every CTLH subunit in our focused sgRNA library.

**Figure 1.**
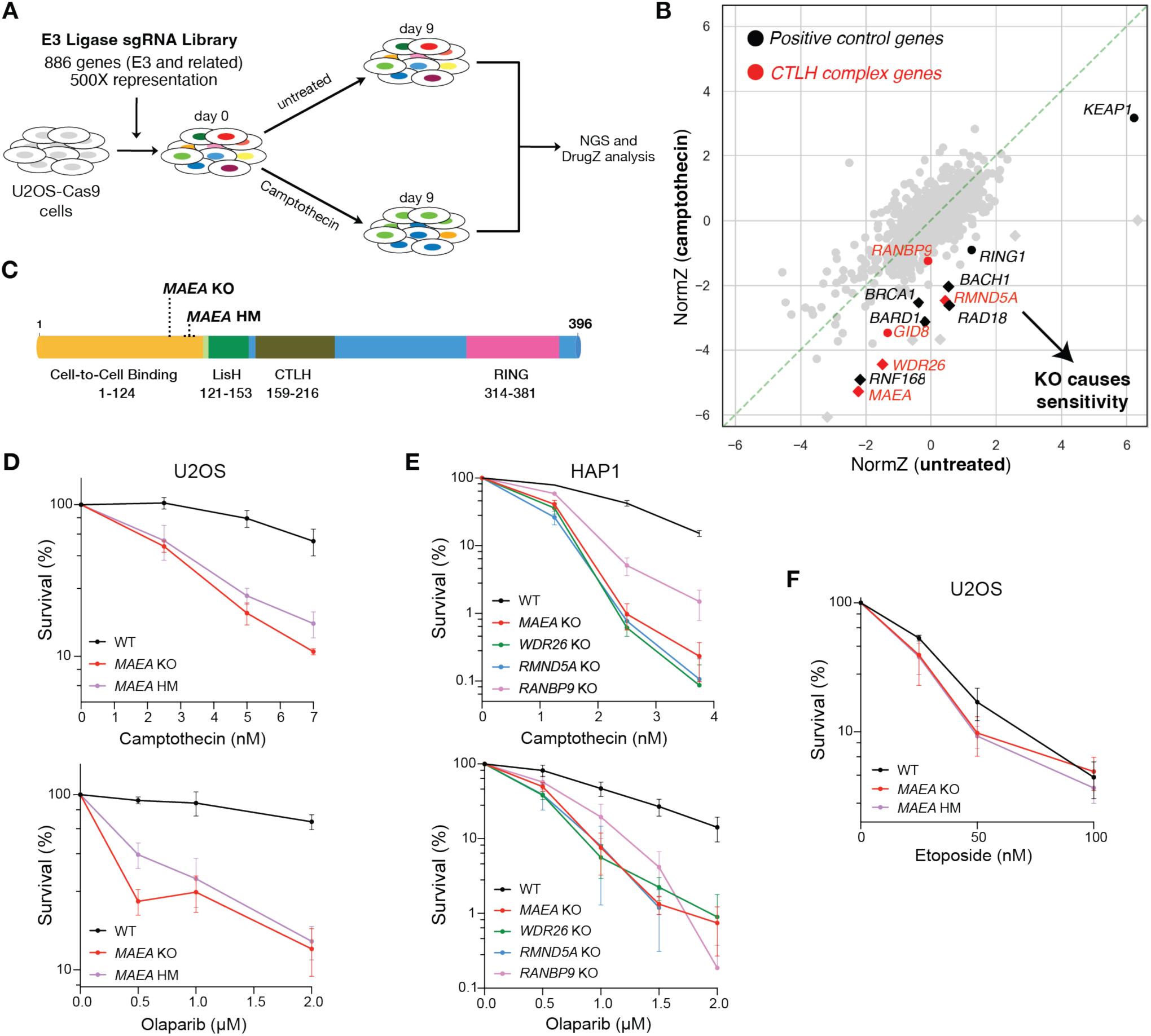
Loss of CTLH components confers hypersensitivity to seDSB-inducing agents. (**A**) Overview of the CRISPR screen. (**B**) DrugZ analysis of (**A**). Black symbols denote positive control proteins, while red symbols denote CTLH complex members. Diamonds and circles denote significant and non-significant hits, respectively. (**C**) MAEA domain map displaying CRISPR-induced edits in *MAEA* KO and *MAEA* hypomorph U2OS cells. (**D**) Clonogenic survival assays in MAEA null U2OS cells with camptothecin and olaparib. (**E**) Clonogenic survival assays in the indicated HAP1 cell lines with camptothecin and olaparib. (**F**) Clonogenic survival assays with etoposide in the U2OS cells lines indicated. Data in (**D-F**) are combined products of three independent experiments. Error bars represent the mean ± SEM. KO = knockout, HM = hypomorph.

While the CTLH complex was known to regulate gluconeogenic enzymes in yeast^18^ and glycolytic enzymes in mammals^19^ our results suggested a previously unrecognized role for it in the DDR. To validate this, we first focused on the key CTLH RING domain subunit MAEA^20,21^. Thus, we generated *MAEA* knockout (KO) and hypomorphic (HM) U2OS cells using CRISPR-Cas9 (**Fig. 1C**, **Supp. Fig. 1A-C)**. The HM allele is a 33**-b**ase pair in-frame deletion leading to loss of amino acid residues 108–117, resulting in a truncated protein that was expressed at lower levels than wild-type (WT) MAEA **(Supp. Fig 1A)**. Compared to WT cells, *MAEA* KO and HM cells were hypersensitive to camptothecin and the PARP inhibitor, olaparib — both of which induce seDSBs during S phase (**Fig. 1D**). Demonstrating that this role was shared with other CTLH subunits and not U2OS-specific, we found that inactivating the genes encoding CTLH components MAEA, WDR26, RMND5A, and RANBP9 led to camptothecin and olaparib hypersensitivity in HAP1 cells (**Fig. 1E**). It was recently shown that HAP1 cells are unusually sensitive to camptothecin due a debilitating mutation in *TDP1*^22^. Further sensitization of HAP1 cells to camptothecin upon CTLH loss suggested that TDP1 and CTLH function in parallel pathways to protect cells from abortive TOP1 lesions.

MAEA loss did not confer detectable hypersensitivity to etoposide (**Fig. 1F**), which induces double-ended DSBs that do not rely much on HR for their repair. Thus, our findings suggested that CTLH may function specifically in HR-related events during S phase. In line with this, we observed that MAEA loss increased the percentage of cells in S phase, supporting the idea that these cells experience replication stress even without exposure to exogenous genotoxic agents (**Supp. Fig. 1D**).

### Clinical *MAEA* variants associated with neurodevelopmental defects in humans hypersensitize cells to seDSB-inducing agents

The CTLH complex plays a role in normal development^23,24^, and while mutations in the core CTLH complex have not been previously associated with pathogenesis, variants in the CTLH substrate adaptor, WDR26, can cause Skraban-Deardorff (SD) syndrome. SD syndrome is characterized by a developmental delay, abnormal gait and seizures^25–30^ and has been associated with compromised CTLH complex assembly and chromatin accessibility^31,32^.

From the Deciphering Developmental Disorders (DDD) database and the 100,000 Genomes Project, and by establishing clinical connections via GeneMatcher, we identified seven individuals with likely pathogenic variants in *MAEA* (**Table 1**, **Supp. Table 7**)^33–37^. All individuals in this cohort presented with developmental delay, intellectual disability, and delayed acquisition of speech. All patients (3 male; 4 female) were between the ages of four and sixteen at the time of evaluation and consistent traits included delayed speech and language acquisition as well as developmental delay and intellectual disability (DD/ID). Most (6/7) had abnormal muscle tone, dysmorphic features, and motor delay. Seizures and autism spectrum disorder were found in a subset (2/7 and 1/7, respectively). In sum, the characteristics observed in these seven patients may constitute a novel nonsyndromic DD/ID. While most individuals (6/7) exhibited some clinical features typically associated with known inherited DNA repair or replication deficiency disorders, others such as seizures, short stature, moderate-to-severe microcephaly, skeletal abnormalities, behavioral issues, immunodeficiency, and bone marrow failure were either absent of displayed only by one or two affected individuals (**Table 1**, **Supp. Table 7**).

**Table 1.** Clinical information of the *MAEA* developmental delay/intellectual disability patient cohort.

| Patient | 1 | 2 | 3 | 4 | 5a | 5b | 6 |
| --- | --- | --- | --- | --- | --- | --- | --- |
| Variant | c.1045G>A | c.900-2A>T | c.816C>A | c.1187T >G | c.1009C>T | c.1009C>T | c. 100C>T |
| Transcript | ENST00000303400.9 | ENST00000303400.9 | ENST00000303400.9 | ENST00000303400.9 | ENST00000303400.9 | ENST00000303400.9 | ENST00000303400.9 |
| Genomic (hg38) | chr4:1338567 | chr4:13332208 | chr4:1336911 | chr4:1339165 | chr4:1338531 | chr4:1338531 | chr4:1312009 |
| Annotation | p.Glu349Lys | Splice site variant before RING | p.Tyr272* | p.Met396Arg | p.Arg337Cys | p.Arg337Cys | p.Arg34Cys |
| Inheritance | De novo | De novo | De novo | De novo | Homozygous | Homozygous | De novo |
| Age at Evaluation | 4 years | 5 years | 9 years 4 months | 13 years 1 month | 8 years | 4 years | 16 years |
| Sex | Male | Male | Female | Female | Female | Female | Male |
| Weight | 14.7 kg (7th centile) | 17.6 kg (25-50th centile) | 30.8kg (75th centile) | 27.2 kg (<%0.4) | 21.5 kg (10-25th centile) | 15.3 kg (10th centile) | 57.3kg (25th-50th) |
| Height | 105 cm (47th centile) | 112 cm (75th centile) | 135cm (91st centile) | 131 cm (<%0.4) | 119 cm (10-25th centile) | Unknown | 174.5cm (50th) |
| OFC | 49.4 cm (25th centile) | Unknown | 51 cm (25-50th centile) | 48.6 cm (age 8y4m; <%0.4) | 45 cm (<2nd centile) | 44 cm (<2nd centile) | 57.3cm (90-97th) |
| Database | GeneMatcher | GeneMatcher | 100,000 Genomes Project | GeneMatcher, DDD | GeneMatcher | GeneMatcher | Cacheiro et al. 2020 |

All but one of the identified patient-associated *MAEA* variants were heterozygous and *de novo*, suggesting that the affected allele provided dominant-negative effects. The other variant (c.1009C>T;p.Arg336Cys), present in two affected siblings (MAEA-P5a and MAEA-P5b) was inherited homozygous and therefore likely recessive. Of note, these siblings also inherited a homozygous variant of uncertain significance (c.317G>A;pArg106His) in *LETM1*, a gene previously linked with a neurodegenerative disorder caused by mitochondrial dysfunction^38^. However, phenotypic comparison of the patients across the cohort coupled with assessments of lactate levels, urine organic acids, and plasma amino acids in patients P5a and P5b indicated that the *LETM1* variant was unlikely to be pathogenic. Moreover, we have identified the same *LETM1* variant in another patient (with no *MAEA* variant) who has no cognitive impairment, further suggesting *LETM1* is unrelated to P5a/P5b’s DD/ID presentations. In summary, the characteristics observed in these seven patients likely constitute a novel form of non-syndromic DD/ID associated with pathogenic variants in *MAEA*, which we have termed DIADEM (**D**evelopmental delay and **I**ntellectual disability **A**ssociated with **DE**fects in **M**AEA).

Further analyses suggested that all patient associated MAEA variants we describe are pathogenic. All missense/truncating MAEA variants except two (those in P2 and P6) are in or near its C-terminal RING domain. The intronic variant present in P2 is predicted to disrupt splicing and create a frameshift mutation that truncates MAEA just before the RING domain. To better understand and predict the impacts of these patient variants, we used existing protein structures^39^ to interrogate the local protein-structural environments of each of the four MAEA patient missense variants within the context of the CTLH catalytic module containing MAEA, RMND5A, UBE2H, and ubiquitin (**Fig. 2A**)^39^. This revealed that Met-396 packs into a hydrophobic pocket on the RMND5A surface that cannot accommodate a substitution with the more bulky and polar arginine side chain found in patient P4. By contrast, Glu-349 engages in electrostatic attraction with a region of positive potential on the ubiquitin surface, and substitution with lysine, as in patient P1, introduces additional steric bulk and apposition of like charges.

**Figure 2.**
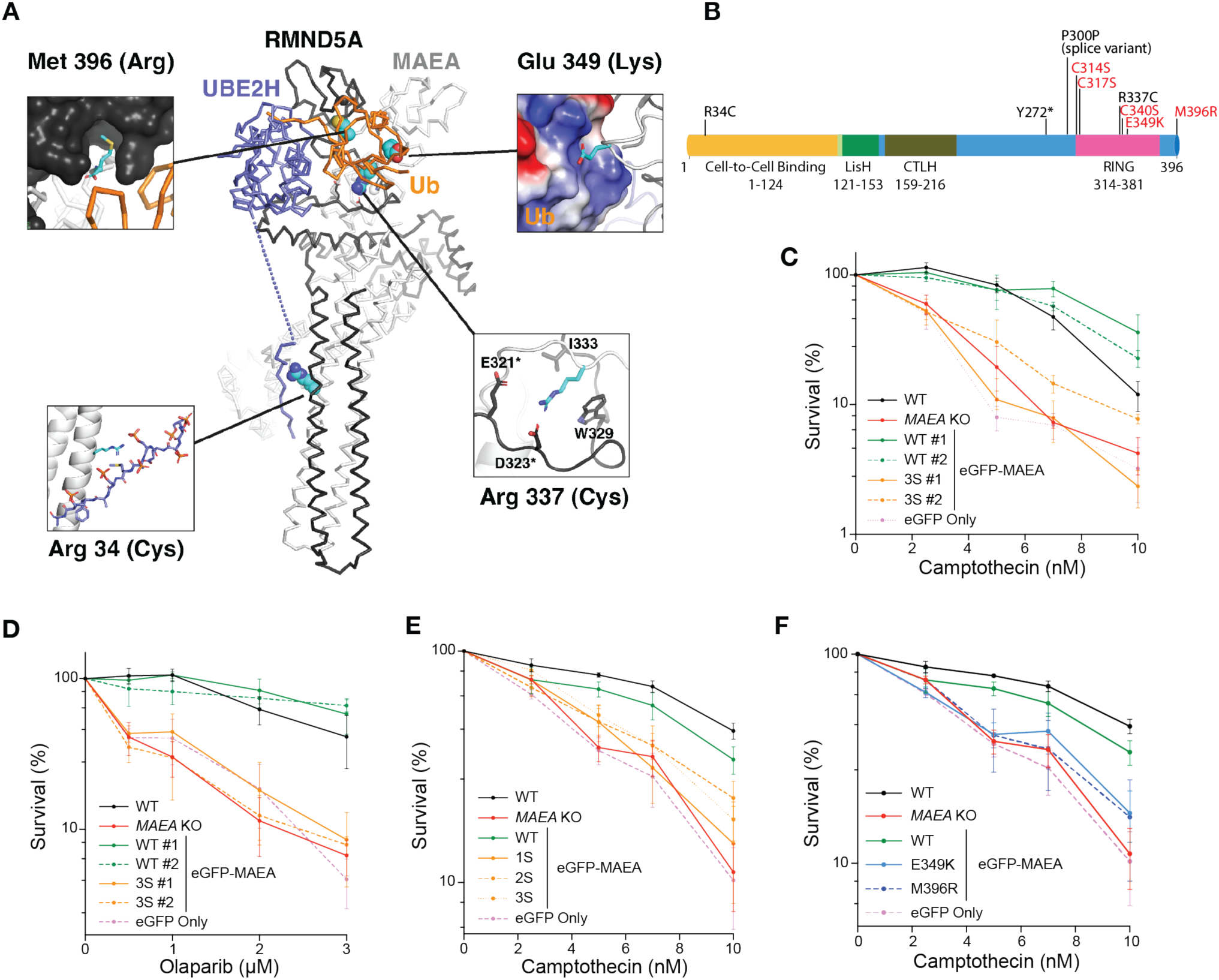
Clinical *MAEA* variants associated with neurodevelopmental defects in humans hypersensitize cells to seDSB-inducing agents. (**A**) Cα trace representation of the MAEA/RMND5A/UBE2H complex crystal structure (PDBID: 8PJN). Altered residues are shown as CPK spheres colored by atom (C-cyan, O – red, N – blue, S – yellow). The environments of each mutant site are shown in the expansion insets. (**B**) Domain map of MAEA with C>S mutations and clinical variants labeled. Bolding indicates cells were complemented with these MAEA variants and tested in subsequent assays. (**C**–**F**) Clonogenic survival assays in the indicated U2OS cell lines with camptothecin or olaparib, as indicated. Data in (**C-F**) are the combined results of 3 independent experiments. Bars represent the mean ± SEM. * P ≤ 0.05, ** P ≤ 0.01, *** P ≤ 0.001, **** P ≤ 0.0001. 1S = eGFP-MAEA^1S^ (C340S), 2S = eGFP-MAEA^2S^ (C314S, C317S), 3S = eGFP-MAEA^3S^ (C314S, C317S, C340S).

Arg-337 engages in both non-polar interactions with tryptophan and valine residues from RMND5A, so replacement with cysteine as in patients P5a and P5b would reduce the extent of these favorable contacts and would also remove other favorable electrostatic interactions with glutamate and aspartate residues from the same RMND5A loop. Finally, the RING-distal substitution of Arg-34 with cysteine in patient P6 appears to directly affect a cluster of phosphorylation-dependent interactions between the MAEA/RMND5A stalk region and the UBE2H C-terminal region that are crucial for formation of the core MAEA/RMND5A/UBE2H complex^40^ (**Fig. 2A**). Together, these analyses suggested that the patient-associated MAEA variants we identified would significantly compromise the integrity of the CTLH ubiquitin ligase complex.

To test whether MAEA ubiquitylation activity is required for its role in the DDR, we complemented *MAEA* KO U2OS cells with various MAEA constructs. These included WT N-terminally tagged eGFP-MAEA (eGFP-MAEA^WT^) and a RING domain mutant (eGFP-MAEA^3S^) version of eGFP-MAEA containing three cysteine to serine (C>S) substitutions: Cys314Ser, Cys317Ser, and Cys340Ser (**Fig. 2B, Supp. Fig. 2A-B**). We mutated these sites based on their evolutionary conservation and their predicted and observed importance for ubiquitylation: Cys314 and Cys317 match the RING domain consensus motif (**C**-X_2_-**C**-X_[9-39]_-**C**-X_[1-3]_-**H**-X_[2-3]_-**C**-X_2_-**C**-X_[4-48]_-**C**-X_2_-**C**) while Cys340 was reported to be essential for MAEA (Gid9) activity in yeast (21). The corresponding cysteine in the RING of RMND5A (Gid2) reportedly plays a similar role (20). We also generated eGFP-MAEA^1S^ (Cys340Ser) and eGFP-MAEA^2S^ (Cys314Ser, Cys317Ser) cell lines for comparison (**Fig. 2B, Supp. Fig. 2B**).

In clonogenic survival assays, expression of eGFP-MAEA^WT^ rescued the hypersensitivity of *MAEA* KO cells to camptothecin and olaparib, while expression of eGFP (hereafter eGFP Only) did not. Furthermore, eGFP-MAEA^1S^, eGFP-MAEA^2S^, and eGFP-MAEA^3S^ variants were expressed at similar levels to eGFP-MAEA^WT^, but failed to complement hypersensitivity to camptothecin and olaparib (**Fig. 2C-E**, **Supp. Fig. 2B**). These investigations thus indicated that the RING domain — and therefore the ubiquitin ligase function — of MAEA is critical for cellular tolerance of seDSB induction.

We next utilised the *MAEA* KO U2OS cells to model the impact of MAEA variants from patients P1 (eGFP-MAEA^E349K^) and P2 (eGFP-MAEA^M396R^) in an isogenic cell background (**Table 1**, **Fig. 2B**). In clonogenic survival assays, unlike eGFP-MAEA^WT^, eGFP-MAEA^E349K^ and eGFP-MAEA^M396R^ failed to complement the camptothecin hypersensitivity of *MAEA* KO cells, despite expressing at similar levels to eGFP-MAEA^WT^ (**Fig. 2F, Supp. Fig. 2B**). This indicated that these clinical variants, like disruption of the MAEA RING domain, confer hypersensitivity to seDSB-inducing agents.

### MAEA loss impairs HR and RAD51 loading but not DNA end resection

To expand on our findings, we used the ‘traffic light reporter’ (TLR) assay to measure HR efficiency^41^ in cells depleted of MAEA or RMND5A. In this system, accurate repair of the chromosomally integrated GFP target only occurs if the induced DSB is repaired by HR with a donor template. As expected, depletion of CtIP, which is crucial for DSB end resection ^42^, greatly impaired HR (**Fig. 3A**). Moreover, following siRNA-mediated depletion of MAEA or RMND5A, we observed a significant reduction (40–80%) of HR relative to cells transfected with a control siRNA against Luciferase (**Fig. 3A, Supp Fig. 3A;** note that the analyses included adjustments for cell cycle profiles). Taken together, these findings indicated a key role for MAEA and the CTLH complex in promoting HR-dependent DNA repair.

**Figure 3.**
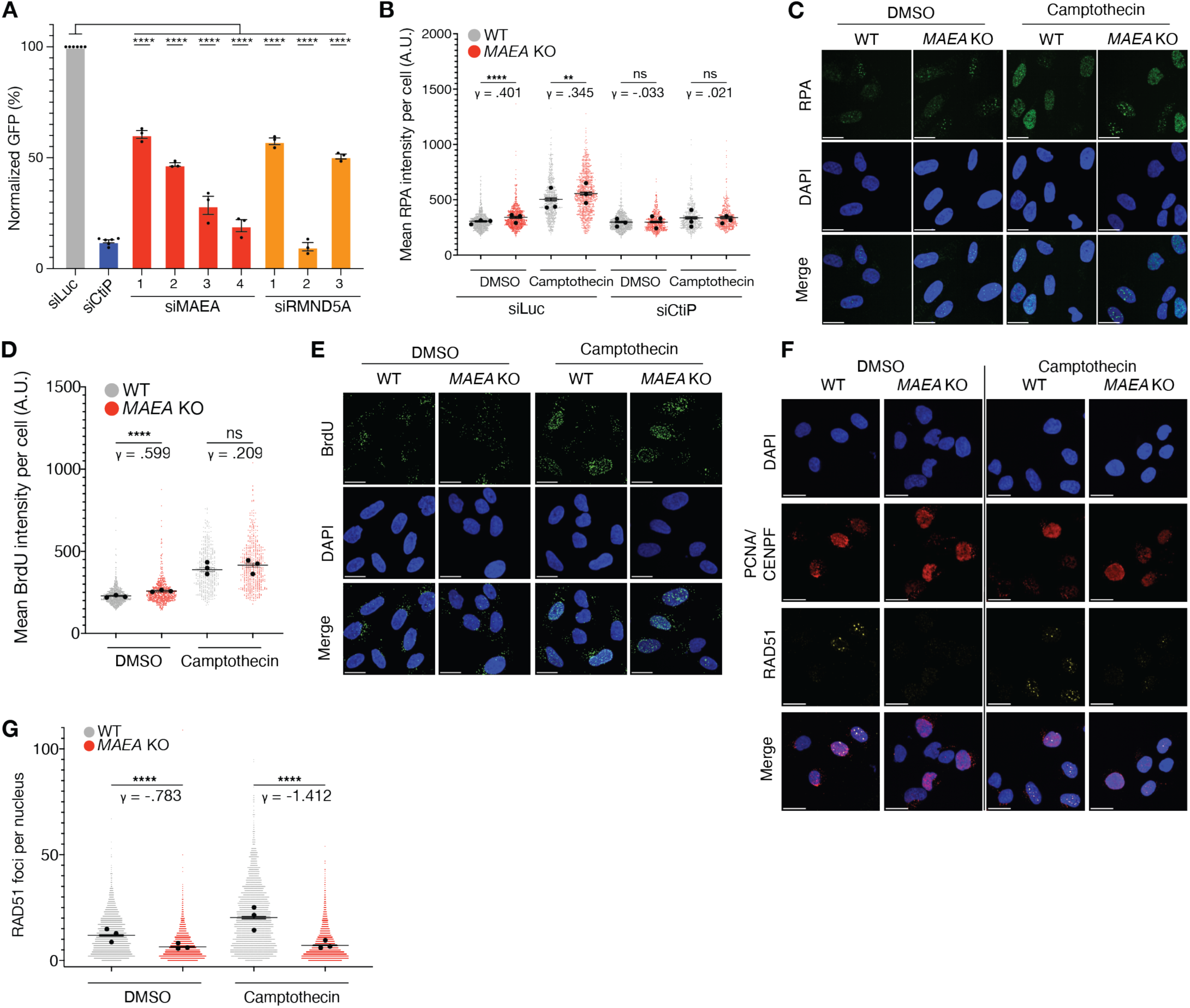
MAEA loss impairs RAD51 loading, but not DNA-end resection. (**A**) TLR assay in U2OS cells with the indicated siRNAs (n = 3). Statistics were generated by performing an ordinary one-way ANOVA comparing each siRNA to siLuc control. siCtIP and siLuc data are from 5 independent experiments. siMAEA and siRMND5A come from 3 independent experiments. (**B-C**) Immunofluorescence-based quantification (**B**) and representative images (**C**) of chromatinised RPA in S phase cells treated with DMSO (1h) or camptothecin (1μM 1h). (**D-E**) Quantification (**D**) and representative images (**E**) of BrdU in U2OS cells treated with DMSO (1h) or camptothecin (1μM 1h). (**F**–**G**) Representative images (**F**) and quantification (**G**) of RAD51 foci in S phase U2OS cells treated with DMSO or camptothecin. All quantifications of foci are the combined results of three independent experiments. Extended fluorescent images from C and E can be found in **Supplemental** Figure 3C-D. For **B**–**H**, P-values were generated by performing a two-tailed Kruskal-Wallis test. γ is a measure of effect size. Data represents three independent experiments. Bars represent the mean ± 95% CI. * P ≤ 0.05, ** P ≤ 0.01, *** P ≤ 0.001, **** P ≤ 0.0001. Scale bars are 20 μm.

We next investigated where CTLH functions in the HR pathway. DNA end resection is a decisive step during DNA DSB processing that dictates its repair by HR mechanisms. Therefore, we examined the levels of single-stranded DNA (ssDNA), as measured by both native BrdU staining and chromatin-bound RPA, in control and *MAEA* null cells following camptothecin exposure^43^. Only cells that were γH2AX positive were included in our analysis, since camptothecin mainly induces DSBs (and therefore γH2AX signal) in S phase. We observed no resection defects in S phase *MAEA* KO cells following camptothecin treatment (**Fig. 3B–E, Supp. Fig. 3B-D**). However, we observed a modest increase in spontaneous native BrdU staining and chromatin-bound RPA levels in undamaged S phase cells lacking MAEA, which was rescued by CtIP depletion (**Fig. 3B–E, Supp. Fig. 3B–D**). These data indicated that MAEA does not hinder camptothecin-induced DSB resection, and suggested that its loss leads to increased resection markers in untreated conditions, which most likely arise from defective HR-dependent resolution of endogenously occurring DNA damage at sites of DNA replication.

Following resection and RPA loading, a key next step in HR is replacement of RPA with RAD51. We thus quantified RAD51 foci formation, a known marker of functional HR^44^, in S phase WT and *MAEA* KO U2OS cells following camptothecin treatment, which both activates HR and induces RAD51-mediated replication fork reversal^2,45^. Compared with WT cells, *MAEA* KO cells exhibited a pronounced failure to form RAD51 foci in S phase cells, despite expressing RAD51 at WT levels (**Fig. 3F-G, Supp. Fig. 3E**). These data indicated that the HR deficiency conferred by MAEA loss is associated with impaired RAD51 loading.

### MAEA loss leads to exacerbated replication stress

Camptothecin and olaparib cause replication stress as well as seDSBs^2^. In unperturbed conditions, compared to WT cells, *MAEA* KO cells were enriched in S/G2 (**Supp. Fig. 1D**), and exhibited elevated ssDNA (**Fig. 3B-E, Supp. Fig. 3B-D**). Since these phenotypes were suggestive of endogenous replication stress, and *MAEA* loss sensitized cells to the ATR inhibitor (ATRi) AZD6738 in a recent study of ours^46^, we hypothesized that MAEA loss might sensitize cells to other replication-stress inducing agents. Thus, we performed clonogenic survival assays with the replication-stress inducing agents hydroxyurea (HU), aphidicolin, and AZD6738. In all cases, we observed hypersensitivity of *MAEA* KO cells to these genotoxins compared with WT cells (**Fig. 4A–C**).

**Figure 4.**
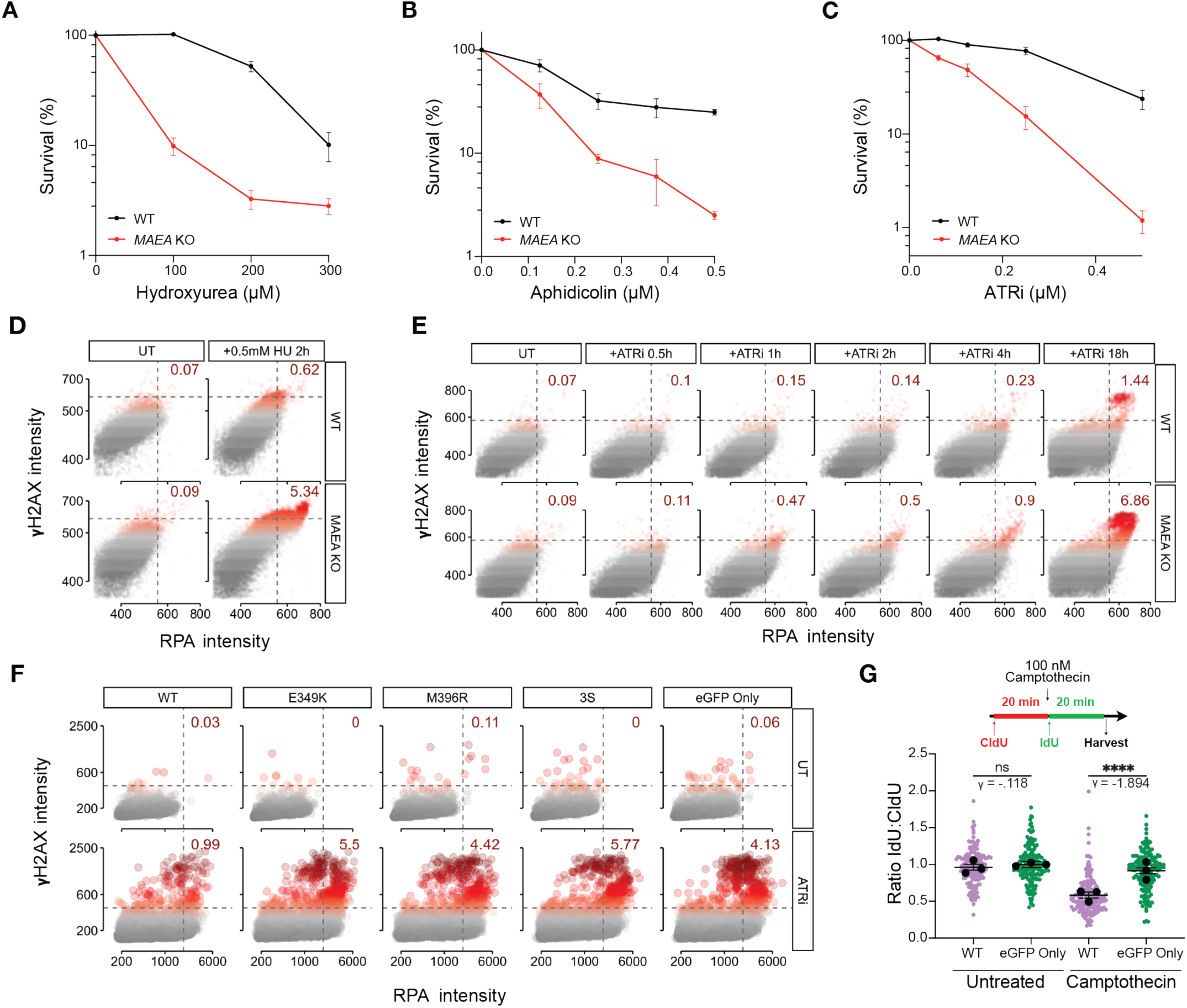
MAEA null cells are hypersensitive to replication stress. (**A**–**C**) Clonogenic survival assays in WT and *MAEA* KO cells with hydroxyurea (HU; **A**), aphidicolin (**B**), and ATRi (**C**). (**D-E**) Quantification of chromatin-associated γH2AX and RPA signal in WT and *MAEA* KO cells upon HU (**D**) or ATRi (**E**) treatment for the indicated times. (**F**) Quantification of chromatin-associated γH2AX and RPA signal upon ATRi treatment in *MAEA* KO U2OS cells complemented with the indicated eGFP expression constructs. Representative images can be found in **Supplemental** Figure 4. (**G**) DNA fiber assay measuring replication fork progression in eGFP Only (*MAEA*^-/-^) and eGFP-MAEA^WT^ cells following treatment with camptothecin. Clonogenic data (**A-C**) are the from 3 independent experiments. Statistics were generated using an ordinary two-way ANOVA. Bars represent the mean ± SEM. DNA fiber data are the combined result of 3 independent experiments. P-values for foci formation and fiber data were generated by performing a two-tailed Kruskal-Wallis test. γ is a measure of effect size. Bars represent the mean ± 95% CI. * P ≤ 0.05, ** P ≤ 0.01, *** P ≤ 0.001, **** P ≤ 0.0001. Scatter plots are representative of two independent experiments. HU = hydroxyurea, ATRi = ATR inhibitor (AZD3768).

These data suggested that MAEA is required for effective HR at damaged replication forks. We investigated this by treating *MAEA* KO and WT U2OS cells with 0.5 mM HU, which primarily induces RAD51-dependent replication fork reversal rather than DSBs^2^. Thus, increased γH2AX signal is likely the result of replication fork collapse. Consequently, we measured the level of γH2AX in cells with replication fork stalling as marked by chromatin-bound RPA. This approach revealed that replicating cells lacking MAEA exhibit increased γH2AX accumulation in *MAEA* KO cells compared with WT cells (**Fig. 4D**). Moreover, *MAEA* KO cells exhibited increased γH2AX accumulation over time compared to WT cells upon ATR inhibition, especially at the later time point of 18 hours (**Fig. 4E**). Using this time point, we measured accumulation of γH2AX in *MAEA* KO cells complemented with eGFP-MAEA^WT^, eGFP-MAEA^E349K^, eGFP-MAEA^M396R^, eGFP-MAEA^3S^, or eGFP Only. In accord with results of clonogenic survival assays with camptothecin and olaparib in these cell lines (**Fig. 2C-F**), we observed an increase in γH2AX levels in the variants and null cell lines compared with eGFP-MAEA^WT^ cells following ATRi treatment (**Fig. 4F, Supp. Fig. 4**). These data implied that the RING domain-dependent E3 ubiquitin ligase function of MAEA is important in managing replication stress and that clinical MAEA variants confer hypersensitivity to replication stress inducing agents. Collectively, our findings indicated that MAEA/CTLH deficiency compromises cellular tolerance of seDSBs and replication stress.

### MAEA protects and promotes DNA replication

To elucidate potential effects of MAEA loss on DNA replication, we utilized DNA fiber spreading assays^47^. We observed a severe replication fork progression defect in the presence of low-dose camptothecin in eGFP Only (*MAEA* KO) cells compared with eGFP-MAEA^WT^ (**Fig. 4G**). This indicated that MAEA loss compromises the ability of cells to resolve replication-stalling TOP1-associated DNA lesions.

We next assessed replication fork dynamics in the absence of exogenous replication stress, and observed that *MAEA* KO cells expressing eGFP Only, eGFP-MAEA^3S^ or two patient-associated variants, eGFP-MAEA^E349K^ and eGFP-MAEA^M396R^, exhibited significantly reduced replication fork progression, associated with increased spontaneous replication fork stalling (**Fig. 5A-B**). These data thus indicated that the E3 ubiquitin ligase activity of the CTLH complex is required to maintain faithful DNA replication in unperturbed conditions.

**Figure 5.**
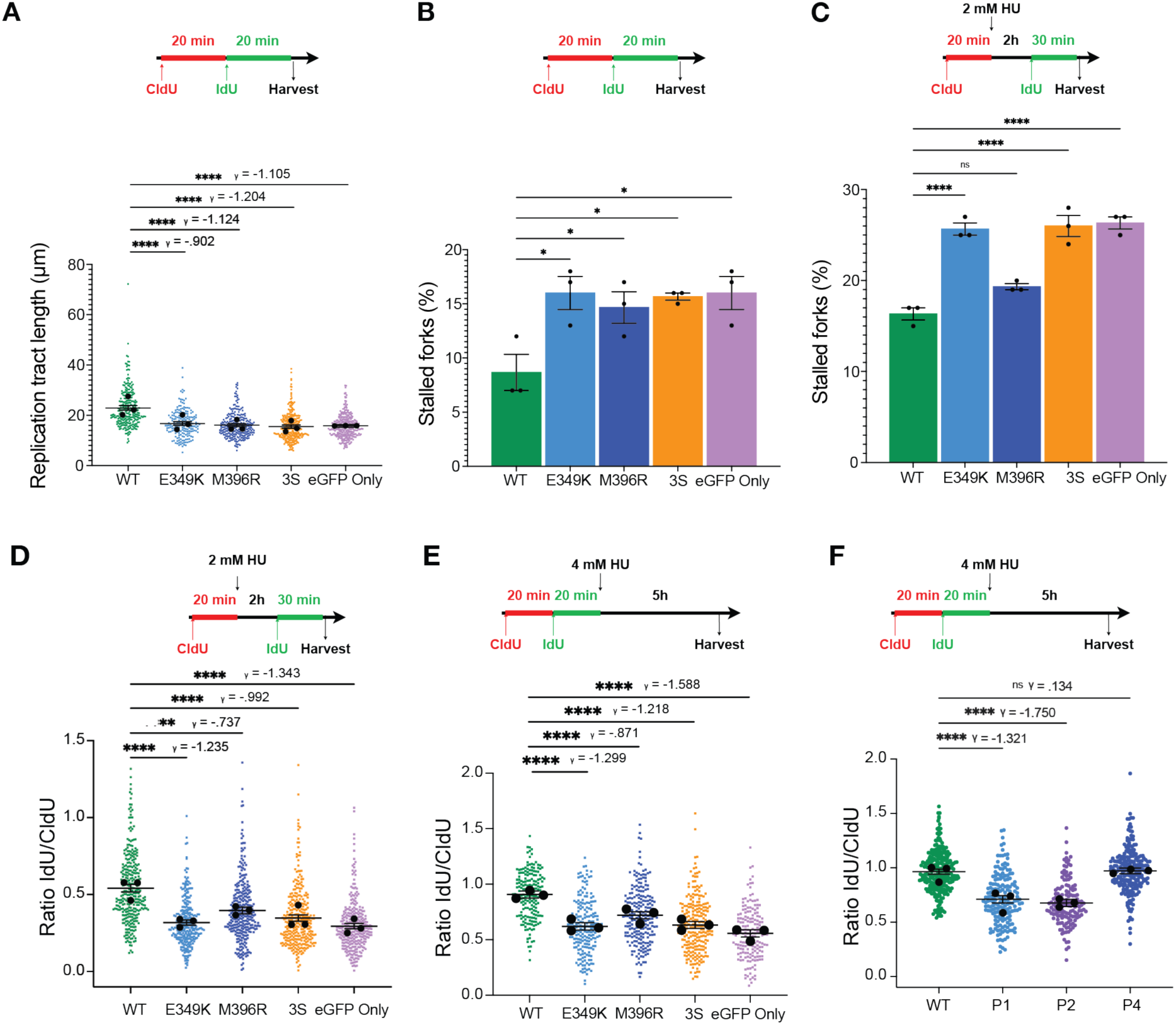
MAEA loss impairs replication fork stability, restart, and protection. (**A-B**) Quantification of replication tract lengths (**A**) and spontaneous replication fork stalling (**B**) in *MAEA* KO U2OS cells complemented with the indicated eGFP expression constructs. (**C-D**) Quantification of HU-induced replication fork stalling (**C**) and fork restart after HU wash-out (**D**) in *MAEA* KO U2OS cells complemented with the indicated eGFP expression constructs. (**E**) Quantification of replication fork degradation after HU treatment in *MAEA* KO U2OS cells complemented with the indicated eGFP expression constructs. (**F**) Quantification of replication fork degradation in primary fibroblasts from Patients 1, 2, and 4 versus a WT control fibroblast cell line following treatment with HU. Data are the combined result of three independent experiments. P-values were generated by performing a two-tailed Kruskal-Wallis test on scatter plots and an ordinary one-way ANOVA on histograms. γ is a measure of effect size. Bars represent the mean ± 95% CI. * P ≤ 0.05, ** P ≤ 0.01, *** P ≤ 0.001, **** P ≤ 0.0001. HU = hydroxyurea.

To assess whether the observed replication fork defects were further impacted by exogenous replication stress, we treated cells with short-term exposure to low millimolar doses of HU, inducing global and synchronous arrest of replication forks that can restart following HU wash-out^2–4^. By conducting DNA fiber assays, we observed that *MAEA* KO cells complemented with eGFP Only or eGFP-MAEA^3S^ exhibited increased fork stalling upon HU treatment and reduced efficiency of fork restart after HU wash-out compared with cells re-expressing eGFP-MAEA^WT^ (**Fig. 5C–D**). This indicated that the E3 ubiquitin ligase function of MAEA is required to properly restart transiently stalled replication forks.

We further observed that while cells expressing eGFP-MAEA^E349K^ exhibited comparable HU-induced replication fork stalling and restart efficiency to *MAEA* null cells, cells expressing eGFP-MAEA^M396R^ exhibited partially reduced replication fork restart efficiency without any apparent increase in replication fork stalling (**Fig. 5C–D**). These observations indicated that while MAEA^E349K^ is likely a null allele, MAEA^M396R^ is a hypomorphic allele that has differential impacts on replication fork stability and restart. In summary, these data indicated that *MAEA* null and variant cell lines struggle with replication fork progression and restart.

RAD51 protects replication forks from nucleolytic degradation during replication fork reversal and aids their remodeling and/or restart^3,48^. Therefore, we considered that increased fork degradation might explain inefficient fork restart in *MAEA* KO cells. Thus, we incubated the panel of *MAEA* KO cells complemented with MAEA variants with CldU and IdU for 20 min each before a 5-hour treatment with 4mM HU to inhibit further DNA replication. We found that MAEA loss or mutation resulted in a reduced IdU:CldU ratio, indicative of increased fork degradation **Fig. 5E**). However, *MAEA* KO cells expressing eGFP-MAEA^M396R^ displayed an intermediate replication fork degradation phenotype, consistent with our other findings assessing replication fork restart (**Fig. 5C–D**) suggesting that this mutation is hypomorphic.

Finally, we established immortalized skin fibroblasts from *MAEA* variant patients based on availability (P1, P2 and P4). Despite still retaining one WT *MAEA* allele, in DNA fiber assays, P1 (p.E349K) and P2 (splice-site mutation prior to the RING domain) cells exhibited significantly enhanced markers of fork degradation after exposure to high dose HU compared with a non-isogenic WT control cell line. Cells from MAEA patient P4 (p.M396R) did not display a fork degradation phenotype, however (**Fig. 5F**). Given that our results modelling this mutation in *MAEA* KO cells indicated that this variant is hypomorphic on a null background (**Fig. 5B–E**), we surmise that in the presence of one WT *MAEA* allele, the impact of the mutant allele is masked.

## Discussion

We have discovered that defects in MAEA and other components of the CTLH ubiquitin E3 ligase complex impair DNA replication and HR, causing hypersensitivity to anti-cancer agents such as hydroxyurea, the PARP inhibitor olaparib, and camptothecin (a close analogue of clinically-used TOP1 poisons). Furthermore, we have identified apparent loss-of-function *MAEA* variants in seven individuals with the nonsyndromic DD/ID that we have termed DIADEM (**D**evelopmental delay and **I**ntellectual disability **A**ssociated with **DE**fects in **M**AEA). All seven DIADEM patients presented with developmental delay, intellectual disability, and delayed acquisition of speech. Additional features including brain abnormalities, reduced muscle tone and unsteady gait, immunodeficiency, and bone marrow failure were also observed with incomplete penetrance. Across the seven identified patients, five bore different *de novo* variants with dominant effects, while the remaining two are siblings (P5a and P5b) and have the same homozygous variant, making it likely that this variant was inherited (**Table 1**).

MAEA is a key subunit of the CTLH E3 ubiquitin ligase complex known to function in development^23,24^, and our findings indicate that several *MAEA* variants likely cause DIADEM by compromising this E3 ubiquitin ligase activity. Notably, loss-of-function variants in another CTLH component (WDR26) causes SD syndrome, which shares some clinical features with DIADEM, but notable differences, including the absence of seizures in five of the patients in our cohort, suggest that DIADEM is distinct from SD syndrome. We speculate that variants in other CTLH components may yield overlapping but non-identical conditions due to their closely connected but non-identical functions.

Our data indicate that cells lacking MAEA are HR deficient, and that while they are proficient in DNA end resection, RAD51 loading onto ssDNA is impaired, a phenotype that can explain the hypersensitivity of these cells towards agents that induce seDSBs. Various core HR factors, including RAD51, BRCA1, and BRCA2, are associated not only with DSB repair but also with cellular tolerance of DNA replication stress^49^. Likewise, MAEA promotes both effective HR and faithful DNA replication. Accordingly, we also found that, in addition to exhibiting increased DSBs and survival defects upon exposure to replication stress-inducing agents, MAEA-deficient cells experience more replication fork stalling under these conditions and less efficient fork restart after release from hydroxyurea.

Based on complementation experiments, we determined that the requirement for MAEA in HR and replication stress depends on its E3 ubiquitin ligase function. In line with our *in silico* analysis of MAEA DIADEM variants, cell models of the patient variants E349K (P1) and M396R (P4) phenocopied hypersensitivity phenotypes observed in the 3S RING domain domain mutant, suggesting that DIADEM can be caused by loss of CTLH ubiquitin ligase activity. However, we note that immortalized fibroblasts from patient P4 did not exhibit detectable replication fork protection defects upon hydroxyurea treatments in contrast to those from patients P1 or P2. Indeed, even in isogenic U2OS cell lines, the M396R variant from patient P4 consistently exhibited milder phenotypes in assays related to replication stress compared with WT MAEA, despite behaving similarly in clonogenic survival assays under camptothecin treatment. We speculate that this variant may have sufficient residual activity to function in some assays. Given the range of variants and the large number of known CTLH substrates^19,50^, the etiology of DIADEM is likely complex and its presentation could depend upon the nature of MAEA variant, as well as the developmental stage of the individual.

While this manuscript was in preparation, another study reported MAEA-mediated polyubiquitylation and degradation of PARP1 in models of gastric cancer and colorectal cancer, with MAEA overexpression conferring cellular sensitivity to oxaliplatin^51^. Given that oxaliplatin induces inter- and intra-strand DNA crosslinks that require HR for their repair^52^, we consider it unlikely that a role for MAEA in regulating PARP1 levels underlies the breadth of HR- and DNA replication-related findings described in our study. Moreover, the study suggests that MAEA loss leads to veliparib (PARP inhibitor) resistance and that, conversely, overexpression generates sensitivity. This is ultimately inconsistent with our own findings that MAEA and CTLH component loss sensitizes cells to PARP inhibition and may be attributable to the specific cell model used in that study.

In our study, MAEA-deficient cells exhibit phenotypes reminiscent of clinical HR deficiency (HRD) as observed in cancer cells. Impaired RAD51 foci formation is a well-established biomarker for HRD and PARPi efficacy in the clinic, making loss or deregulation of the CTLH complex potential prognostic biomarkers for PARPi treatments. Several CTLH components have been found in CRISPR screens as candidate mediators of PARP inhibition and replication stress^46,53–56^, but until now these phenotypes have remained unexplored. Highlighting the potential for CTLH components as prognostic biomarkers, we note that loss of expression of CTLH components *MAEA* and *YPEL5* have been reported in subsets of tumor types (e.g., breast, ovarian) for which PARPi therapies are approved^56^.

Unlike *BRCA1* or *BRCA2* deficiency, DIADEM does not currently appear to predispose to cancer. However, due the young age of the patients in our cohort, the impact of MAEA variants on ageing-associated diseases is unknown; long-term monitoring of these patients will be an important aspect of their care. Moreover, based on our findings, we caution that should a patient with DIADEM develop cancer, the use of chemotherapies that are highly effective in HRD cancers (platinum agents, olaparib, and derivatives of camptothecin such as irinotecan and topotecan) might lead to severe toxicity. While not the focus of our study, such considerations might be relevant to patients with pathological variants in other CTLH complex members, including SD syndrome patients.

## Materials and Methods

### Study approval

Written informed consent to publish clinical information and photographs of the affected individuals was obtained from the families prior to their involvement in this study, in accordance with local IRB-approved protocols. Further approval for this research was obtained from the West Midlands, Coventry, and Warwickshire Research Ethics Committee (Coventry, United Kingdom; REC: 20/WM/0098).

### CRISPR-Cas9 Screen

A custom library of CRISPR guides (**Supplemental Table 4**) targeting 886 E3 ligases and related proteins was cloned into pKLV2-U6gRNA5(BbsI)-PGKpuro2ABFP-W (Addgene Plasmid #67974), packaged into lentiviral particles using second generation plasmids psPax2 (Addgene Plasmid #12260) and pMD2.G (Addgene Plasmid #12259), and titered. Three independent populations of WT U2OS-Cas9 cells were transduced at a multiplicity of infection of 0.25 at 500X representation and selected using puromycin (2μg/mL) for 14 days. Cells were subjected to IC_50_ camptothecin (12nM, 9 days) or no treatment. Both treated and untreated cells were allowed to recover for three additional days with no treatment. Throughout, cells were cultured in Dulbecco’s Modified Eagle Medium (DMEM) with 1X Penicillin-Streptomycin-Glutamine (PSQ, Thermo, Catalog #10378016) and 10% fetal bovine serum (FBS), as well as 10 μg/mL BSD to select for Cas9 expression. DNA from pre- and post-treatment pooled cell populations were extracted and PCR amplified with relevant next-generation sequencing barcodes. Sequencing results were analyzed by using DrugZ^57^.

### Cell culture and generation of cell lines

U2OS Cas9 cells were transfected with one of two guides (IDT) targeting MAEA, designed using Guide Picker and CRISPOR (**Supplemental Table 2**)^58,59^. After transfection with RNAiMAX (Thermo Fisher Scientific), cells were diluted and grown as single colonies. Genomic DNA extraction, PCR, Sanger sequencing and Tracking of Indels by DEcomposition (TIDE) analysis^60^ was used to identify candidate KO clones. Dermal primary fibroblasts were grown from skin-punch biopsies and maintained in DMEM supplemented with 20% FCS, 5% l-glutamine, and 5% penicillin-streptomycin antibiotics (Merck). Primary fibroblasts were immortalized by lentiviral transduction with hTERT that was generated by transfecting 293FT cells (Thermo Fisher Scientific) with pLV-hTERT-IRES-hygro (Addgene Plasmid #85140) and second-generation lentivirus packaging plasmids as above. Cells were selected with hygromycin (Thermo Fisher Scientific) at 70 μg/mL. U2OS, HAP1, and RPE-1 cells were grown at 37°C in 5% CO_2_. U2OS cells were grown in DMEM supplemented with 1X Penicillin-Streptomycin-Glutamine (PSQ, Thermo, Catalog #10378016) and 10% fetal bovine serum (FBS). RPE-1 cells were maintained in DMEM/F-12 medium supplemented with 1X PSQ and 10% FBS. HAP1 cells were maintained in Iscove’s Modified Dulbecco’s Media (IMDM) supplemented with 1X PSQ and 10% FBS. Trypsin-EDTA (Gibco, Trypsin-EDTA (0.25%), phenol red, Catalog #25200056) was used for cell passaging except where otherwise indicated.

### hTERT fibroblast immortalization

Dermal primary fibroblasts were grown from skin-punch biopsies and maintained in Dulbecco’s modified Eagle’s medium (DMEM; Thermo Fisher Scientific) supplemented with 20% FCS, 5% L-glutamine and 5% penicillin-streptomycin. Primary fibroblasts were immortalized with a lentivirus expressing human telomerase reverse transcriptase (hTERT) that was generated by transfecting 293FT cells (Thermo Fisher Scientific) with the plasmids: pLV-hTERT-IRES-hygro (Addgene Plasmid #85140), psPax2 (Addgene Plasmid #12260) and pMD2.G (Addgene Plasmid #12259). Selection was performed using hygromycin (Thermo Fisher Scientific) at 70 μg/mL.

### Clonogenic survival assays

6-well plates were seeded with 500 U2OS or 250 HAP1 cells in technical triplicate. After 8h or overnight of attaching/growth following initial seeding, all medium was replaced with medium containing the appropriate drug dilution. After 10-14 days, cells were washed with PBS and stained with crystal violet. Colonies were counted after scanning plates and using FIJI. DNA damaging agents: olaparib (.5–2μM U2OS), etoposide (25–100nM U2OS), camptothecin (2.5–7 nM U2OS). Data shown represent a minimum of 3 independent experiments. Each experiment consists of the averaged values from 3 technical replicates.

### Immunoblotting

Following SDS-PAGE and wet transfer, PVDF membranes (Amersham Hybond P 0.45) were blocked in 5% BSA/TBST for 1 h at room temperature on rocker. Membranes were incubated with primary antibodies (**Supplemental Table 5**) for 1 h at room temperature, washed 3 times, and then incubated for 45 min with the HRP-conjugated secondary antibody at room temperature. Signals were detected using chemiluminescence and an X-ray film processor.

### Immunofluorescence

Cells were seeded at ∼40K per well in 24-well glass-bottom plate. Where appropriate, cells were grown with BrdU (Merck, B9285). After 24 h, cells were treated with CPT (camptothecin; 1 μM, 1 h) or DMSO. Cells were washed and pre-extracted with ice cold CSK (25 mM HEPES pH 7.4, 50 mM NaCl, 3 mM MgCl_2_, .5% Triton X-100, 300 mM sucrose, ddH_2_O) on ice (3 min) prior to fixation in 2% PFA (10 min). Cells were washed with PBS-0.2% Tween20 (PBST), blocked in 5% BSA and stained with primary antibodies (see **Supplemental Table 6** for details). Cells were washed with PBST and incubated in the dark with secondary antibodies, and 2 μg/mL DAPI. Images were acquired using an Opera Phenix Plus microscope and analyzed with Harmony v.5.1 software. Cells were S phase-stratified using cell cycle PCNA/CENPF or γH2AX.

### Traffic light reporter (TLR) assay

TLR assays were conducted as described previously^12^. Cells were counted on either a Beckman CytoFLEX LX or Thermo Fisher Attune NxT (consistent within set of replicates). Data was analyzed on FlowJo. Gates were set with an internal negative control population expressing neither BFP nor IFP. Background cell counts were subtracted from all totals. Data represent at least three independent experiments. siLuciferase, a negative control, and siCtIP, targeting a known HR factor, were performed six times.

### Statistical tests

The unpaired t-test, two-tailed Kruskal-Wallis test, and ordinary one-way Analysis of Variance (ANOVA) tests were generated using GraphPad Prism v.10.01. Effect size (γ) using the C# GitHub repository Perfolizer by Andrey Akinshin (https://github.com/AndreyAkinshin/perfolizer)^61^. γ is similar to a non-parametric Cohen’s d and is a measure of the pooled median absolute deviations that fit between the median of group X (e.g. *MAEA* WT) and the median of group Y (e.g. *MAEA* KO). In general, the absolute value of γ can be evaluated according to the guidelines 0.00 < 0.10 (Negligible), 0.10 < 0.20 (Weak), 0.20 < 0.40 (Moderate), 0.40 < 0.60 (Moderately Strong), 0.60 < 0.80 (Strong), and 0.80 < (Very Strong).

### DNA fiber assay

Cells were pulse-labeled with 25 μM CldU for 20 min, washed with PBS, pulse-labeled for 20 min with 250 μM IdU, and then harvested. For replication restart experiments, cells were labeled with CldU for 20 min, washed in warm PBS, and incubated in medium containing 2 mM HU for 2 h. Cells were washed again in warm PBS and then incubated with 250 µM IdU for 20 min. Replication in the presence of replication stress was assayed by first pulse labeling cells with 25 µM CldU, which was washed off with medium containing 250 µM IdU and 100 nM CPT. Cells were then pulse labeled with 250 µM IdU and 100 nM CPT, for untreated cells CPT was omitted, at the end of the pulse labeling the cells were harvested as previously described. For fork protection experiments, cells were labeled with 25 μM CldU for 20 min, washed with CldU containing medium, labeled with 250 μM IdU for 20 min, washed with warmed medium containing 4 mM HU, and incubated in medium containing HU for 5 h. DNA fiber analysis was carried out as previously described^62^.

### Structural analysis

The potential impact of the MAEA mutations was assessed by inspection of the molecular context of each variant residue in the MAEA/RMND5A/UBE2H complex crystal structure (PDBID: 8PJN^39^) using PyMOL (PyMOL molecular graphics system, V 3.0, Schrodinger LLC). Missing side chains were modelled manually by selecting rotamers with the lowest clash score.

## Acknowledgements

We thank the patients and parents for taking part in this study and generously donating tissue samples. We thank M. Agudo for help with the CRISPR screen. We acknowledge K. Harnish of the Gurdon Institute for performing next-generation sequencing of our CRISPR-Cas9 screening samples and the Gurdon and CRUK Cambridge Institute core facilities for assistance and support. Research in the SPJ laboratory is supported by Cancer Research UK (CRUK) Discovery Award DRCPGM\100005, CRUK Cambridge Institute core grant SEBINT-2024/100003 and ERC Synergy Award 855741 (DDREAMM). SH was supported by Wellcome Investigator Award 206388/Z/17/Z and CRUK Discovery Award DRCPGM\100005; SWA by a Mark Foundation for Cancer Research (MFCR) ASPIRE II Award; is a recipient of the Women’s Postdoctoral Career Development Award in Science from the Weizmann Institute of Science; and was a recipient of an Outstanding Postdoctoral Women Fellowship from the Israeli Council for Higher Education; SL by ERC Synergy Award 855741; JCT by Wellcome Investigator Award 206388/Z/17/Z and ERC Synergy Award 855741; CJC, GZV, RB and YG by CRUK Discovery Award DRCPGM\100005; ASB by CRUK RadNET Cambridge C17918/A28870 and Wellcome Early Career Award 227014/Z/23/Z; and OL by CRUK Cambridge Institute core grants C9545/A29580 and SEBINT-2024/100003. The SPJ laboratory was also supported by CRUK Programme grant C6/A18796 and core funding grants C6946/A24843 and WT203144 to the Gurdon Institute. GSS and SSJ are funded by a CRUK Programme grant (C17183/A23303) and an MRC project grant (UKRI577). CJC is supported by a grant from the Deutsche Forschungsgemeinschaft (DFG; KU 563/18-1). Research in the Beli lab is funded by the Deutsche Forschungsgemeinschaft (German Research Foundation, DFG) project-ID 393547839—SFB 1361. This research was made possible through access to data in the National Genomic Research Library, which is managed by Genomics England Limited (a wholly owned company of the Department of Health and Social Care). The National Genomic Research Library holds data provided by patients and collected by the NHS as part of their care and data collected as part of their participation in research. The National Genomic Research Library is funded by the National Institute for Health Research and NHS England. The Wellcome Trust, Cancer Research UK and the Medical Research Council have also funded research infrastructure. For the purpose of open access, we have applied a Creative Commons Attribution (CC BY) public copyright licence to any Author Accepted Manuscript version arising from this submission.

## Author Contributions

SHH, YG, and SPJ conceived the project. SHH carried out CRISPR screens, clonogenic survival assays, and other experiments, as well as drafting the manuscript. SWA carried out replication stress assays. GSS obtained patient samples and generated patient-derived cell lines. SSJ conducted DNA fiber assays and analysis. SPJ, GSS, YG, CJC and GZV supervised the project. SJS provided structural analysis of patient variants. LB, SM, SWA, EC, ND, EC, PC, AM, DS, and IT provided clinical insights from their patients. TM and OL provided experimental support. PB supervised TM. RB provided support in TLR assay and supervised OL. AB, SL, and JCT provided bioinformatic and statistical support. All authors contributed to manuscript preparation.

---

**Supplemental Figure 1.**
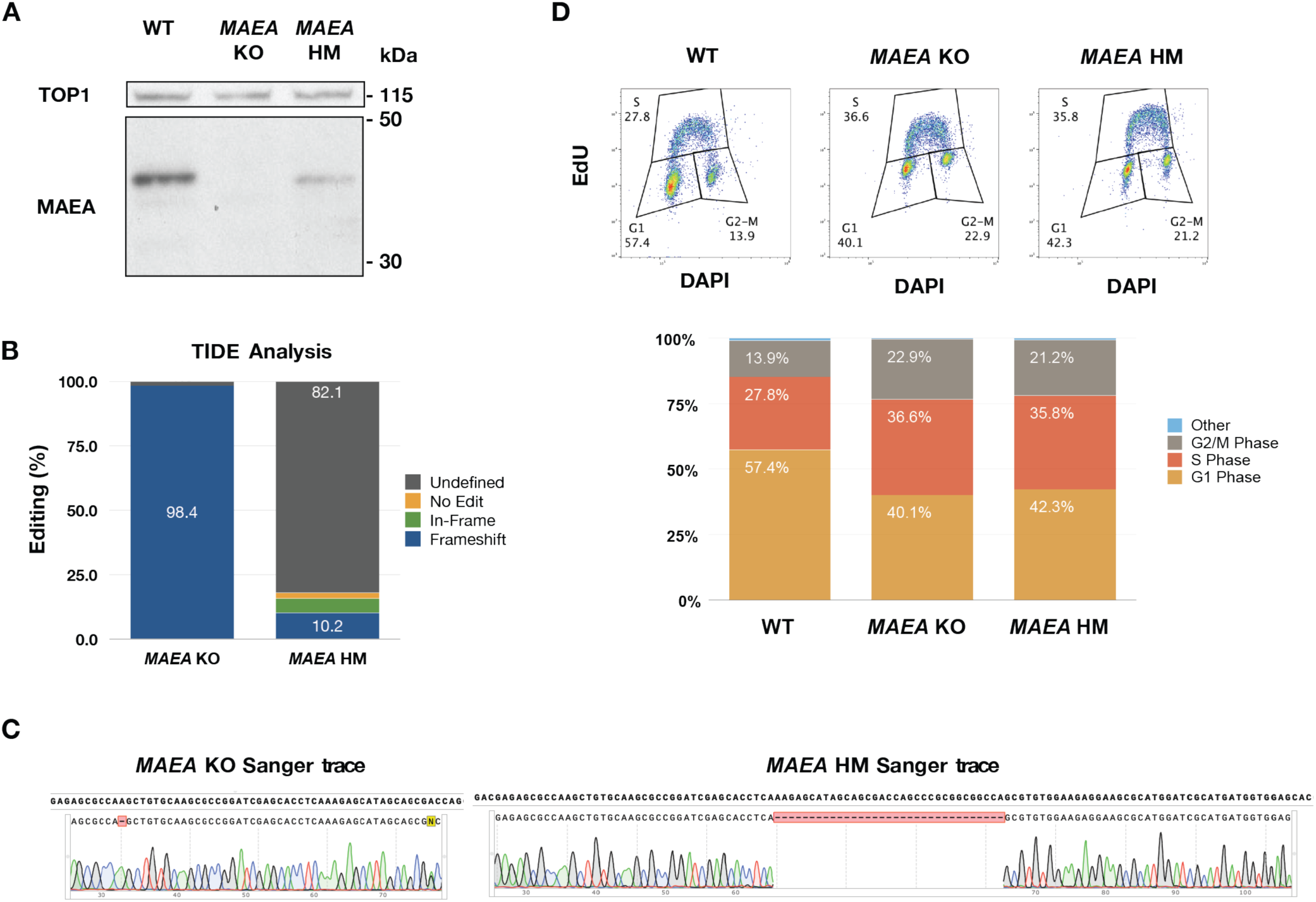
M*A*EA KO and HM cell line validation. (**A**) Immunoblot for MAEA in the indicated U2OS cell lines. (**B**) TIDE analysis of the sgRNA target site in *MAEA* KO and *MAEA* HM cells. (**C**) Sanger trace analysis of the sgRNA target site in *MAEA* KO and HM cells. (**D**) Cell cycle analysis of *MAEA* KO and HM cells.

**Supplemental Figure 2.**
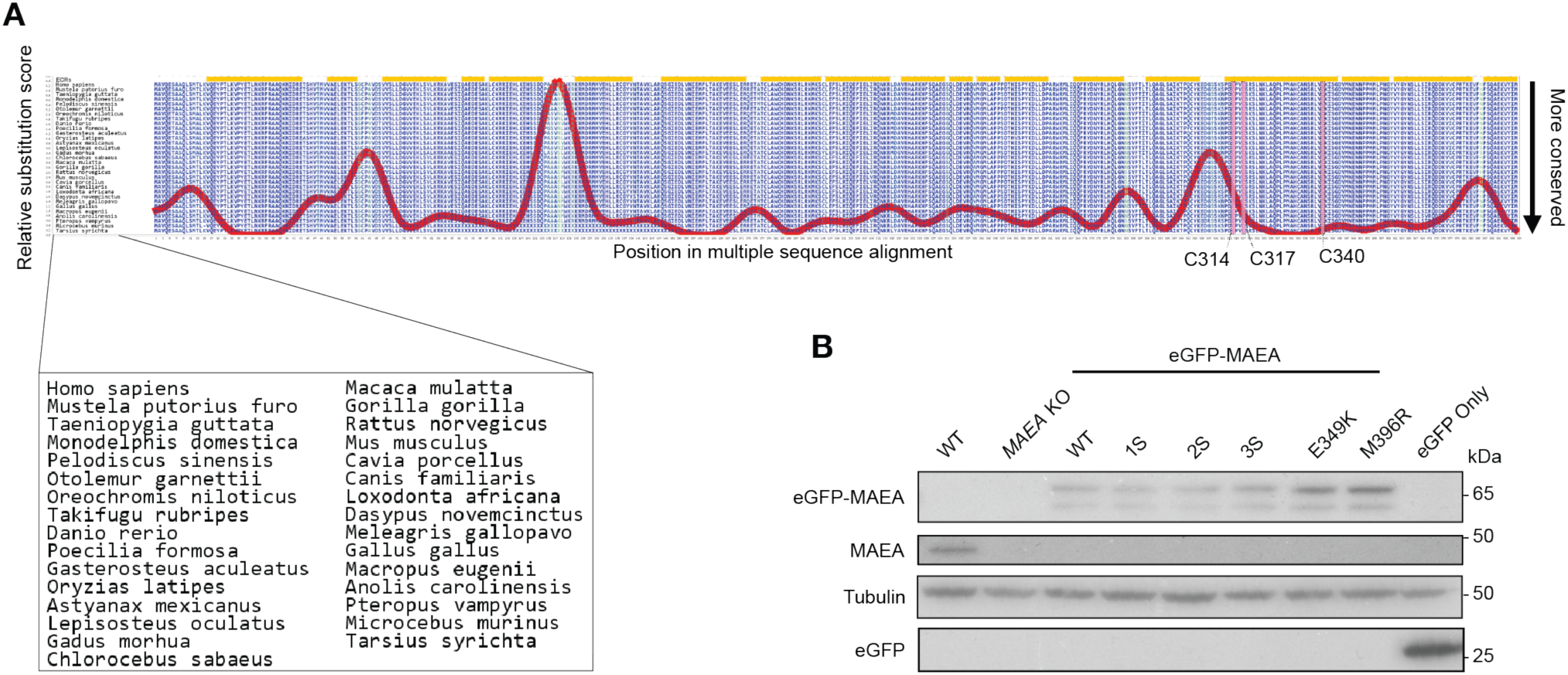
MAEA loss compromises RAD51 foci formation. (**A**) Immunoblot against the indicated proteins with the indicated siRNAs. (**B**) Quantification of BrdU immunofluorescence, after camptothecin treatment in the presence or absence of siRNA-mediated CtIP depletion. (**C**–**D**) Representative images from **Fig. 2C,E** expanded to include γH2AX staining, indicative of S phase cells. (**E**) Immunoblot against RAD51 in WT and *MAEA* KO U2OS cells. P-values were generated by performing a two-tailed Kruskal-Wallis test. γ is a measure of effect size. Data are the combined results of two independent experiments. Bars represent the mean ± 95% CI. * P ≤ 0.05, ** P ≤ 0.01, *** P ≤ 0.001, **** P ≤ 0.0001. Scale bars are 20 μm. NT = non-treated.

**Supplemental Figure 3.**
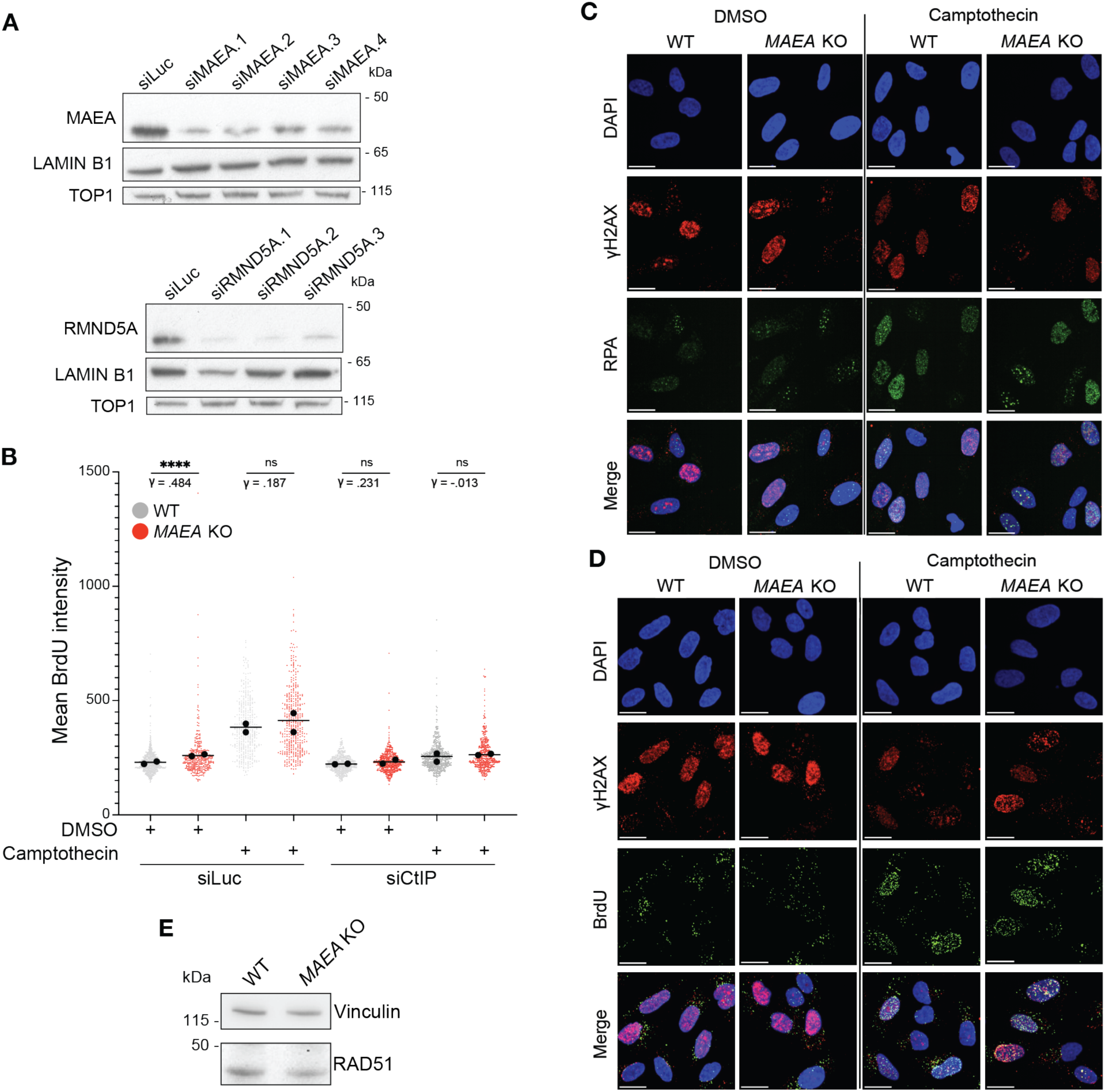
Clinical and C>S MAEA variants are evolutionarily conserved. (**A**) Aminode^63^ analysis of MAEA, with C>S mutations annotated. The red line represents conservation across species. Species compared in analysis are listed in box. (**B**) Immunoblot assessing expression of eGFP-MAEA constructs.

**Supplemental Figure 4.**
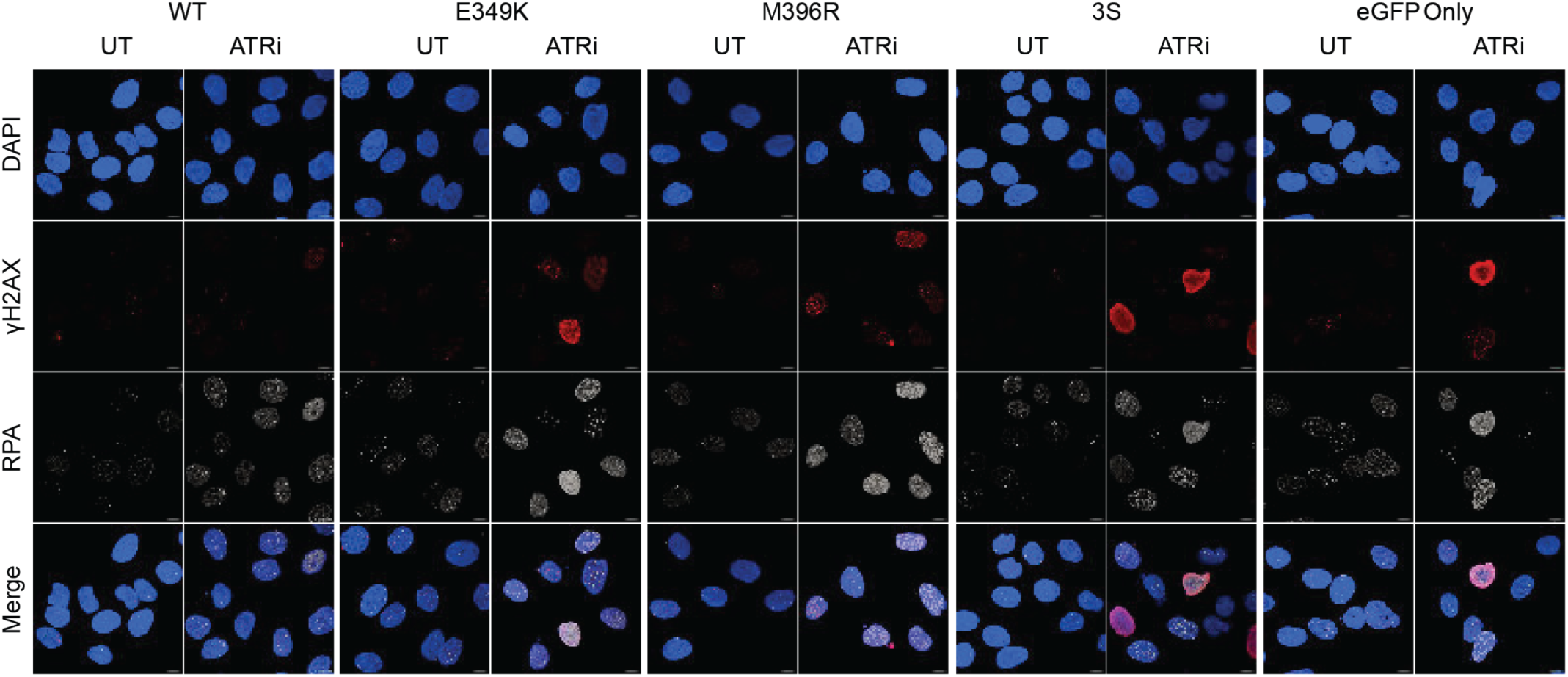
Clinical MAEA variants and loss compromise genome integrity upon ATR inhibition. Representative confocal images from **Fig. 5F**.

